# Horizontal transfer and finalization of a reliable detection method for the olive fruit fly endosymbiont, *Candidatus* Erwinia dacicolax

**DOI:** 10.1101/326090

**Authors:** Gaia Bigiotti, Roberta Pastorelli, Roberto Guidi, Antonio Belcari, Patrizia Sacchetti

## Abstract

**Background:** The olive fly, *Bactrocera oleae*, is the most important insect pest in olive production, causing economic damage to olive crops worldwide. In addition to extensive research on *B. oleae* control methods, scientists have devoted much effort in the last century to understanding olive fly endosymbiosis with a bacterium eventually identified as *Candidatus* Erwinia dacicola. This bacterium plays a relevant role in olive fly fitness. It is vertically transmitted, and it benefits both larvae and adults in wild populations; however, the endosymbiont is not present in lab colonies, probably due to the antibiotics and preservatives required for the preparation of artificial diets. Endosymbiont transfer from wild *B. oleae* populations to laboratory-reared ones allows olive fly mass-rearing, thus producing more competitive flies for future Sterile Insect Technique (SIT) applications.

**Results:** We tested the hypothesis that *Ca.* E. dacicola might be transmitted from wild, naturally symbiotic adults to laboratory-reared flies. Several trials have been performed with different contamination sources of *Ca.* E. dacicola, such as ripe olives and gelled water contaminated by wild flies, wax domes containing eggs laid by wild females, cages dirtied by faeces dropped by wild flies and matings between lab and wild adults. PCR-DGGE, performed with the primer set 63F-GC/518R, demonstrated that the transfer of the endosymbiont from wild flies to lab-reared ones occurred only in the case of cohabitation.

**Conclusions:** Cohabitation of symbiotic wild flies and non-symbiotic lab flies allows the transfer of *Ca.* E. dacicola through adults. Moreover, PCR-DGGE performed with the primer set 63F-GC/518R was shown to be a consistent method for screening *Ca.* E. dacicola, also showing the potential to distinguish between the two haplotypes (htA and htB). This study represents the first successful attempt at horizontal transfer of *Ca.* E. dacicola and the first step in acquiring a better understanding of the endosymbiont physiology and its relationship with the olive fly. Our research also represents a starting point for the development of a laboratory symbiotic olive fly colony, improving perspectives for future applications of the Sterile Insect Technique.

## Background

Relationships between fruit flies (Diptera: Tephritidae) and microorganisms, especially bacteria, have been studied for a long time. Much research has focused on the biology and behaviour of many of these flies, but their symbiotic associations have been less investigated. In particular, the role that these microorganisms could play in fly biology, physiology and behaviour has not been well studied [1, 2]. One of the most questionable issues in this research area, which scientists are still working on, is the relationship between the olive fruit fly *Bactrocera oleae* (Rossi) and its associated bacteria [3, 4, 5, 6]. In particular, symbiotic bacteria seem to be necessary for this Tephritid’s fitness [7, 8]. Furthermore, recent studies have shown that symbiosis plays a very relevant role in the *B. oleae*’s lifespan [9, 10]. Thus, symbiosis in the olive fruit fly is considered to be very important to understand its behaviour and its life cycle.

At the beginning of the 20th century, Petri [11] was the first scientist who described the bacteria inside the *B. oleae* gut, both in larvae and adults; later, other scientists tried to better define this endosymbiosis [5, 8, 12, 13]. More recently, thanks to the advent of biological molecular techniques, such as PCR amplification and sequencing, the *B. oleae* endosymbiont was identified as *Candidatus* Erwinia dacicola [14]. It was found only in wild *B. oleae* flies, and even if it could not be cultivated, it was supposed to be more abundant than other bacteria. It was therefore assumed to be a tightly associated endosymbiont of the olive fruit fly [15]. *Ca.* E. dacicola lives and multiplies inside a small organ of olive fruit flies, which Petri first described as a “cephalic vesicle” or “pharyngeal gland” [11]. In more recent studies, this organ is referred to as the “oesophageal bulb” [6, 14, 16]. Despite this, the symbiont has been detected in other adult organs, including the gut and the last digestive tract near the ovipositor [14, 15].

*Ca.* E. dacicola has been assigned to the Enterobacteriaceae family within the γ-Proteobacteria group [14] and is considered a P-symbiont (persistent) for *B. oleae.* It is vertically transmitted through generations, from the female to the egg, and it has been found in every stage of the fly lifespan, particularly in the adult one. In addition, it was shown that *Ca.* E. dacicola seems to switch from an intracellular existence to an extracellular one during the host insect development, since it lives intracellularly within cells of the larval midgut caeca and extracellularly in the adult gut [15]. Recent studies have highlighted the fact that the larvae can develop in unripe olives, owing to the presence of *Ca.* E. dacicola presence [17]. According to this, the endosymbiont strictly affects the larval survival of unripe olives. Larvae, thanks to *Ca.* E. dacicola, are able to overcome the effects of some compounds such as oleuropein, which seems to be detrimental, acting as an anti-nutrient and allowing both larval development and a higher nitrogen level assumption. Along with this, oleuropein may inactivate enzymes or reduce the digestibility of dietary proteins, preventing larvae from assuming nutrients [17].

The symbiont seems to be strictly related to the olive tree agro-ecosystem, since its presence has never been confirmed in laboratory-reared flies [6, 10] with the exception of a recent research in which the bacterium was found in few specimens of a lab hybrid population [18].

*B. oleae* is a fruit fly that is difficult to rear artificially; however, long lasting research has demonstrated that there are still several mass rearing difficulties, including high costs and labourintensive procedures [19]. Lab colonies are usually obtained from lab-adapted wild populations. Flies often will not easily oviposit in artificial rearing devices such as wax domes and tend not to develop well on a cellulose-based artificial diet, two essential aspects of the mass rearing technique [20]. Previously, when *B. oleae* was reared through these procedures for a long time, several genetic and biological changes appeared [21] as well as behavioural modifications [22]. This suggests that an endosymbiont lacking in lab-reared flies could be involved in all these rearing issues. The absence of *Ca.* E. dacicola in lab-reared colonies could also be caused by the widespread use of antibiotics in the artificial diet; importantly, recent studies have demonstrated that *B. oleae* can be reared without antibiotics [23]. In this way, the endosymbiont might not be lost.

To improve mass-rearing and to produce more competitive flies, it would be favourable to transfer the endosymbiont from wild *B. oleae* populations into lab reared flies in order to start up Sterile Insect Technique (SIT) field applications. This would allow for the release of sterile and more competitive males due to the endosymbiont *Ca.* E. dacicola. This would likely be a more effective and highly sustainable method to reduce *B. oleae* field populations.

Moreover, recent research has highlighted the endosymbiont presence in reared flies, demonstrating that the endosymbiont may have entered the lab colony during cohabitation with wild flies [18]. Along with horizontal transfer, it is important to determine the precision and reliability of the *Ca.* E. dacicola DNA detection procedure. Since 2005, endosymbiont presence has been detected many times in wild flies, both in larvae and adults. However, its DNA has never been confirmed using the same set of primers [6, 14, 15, 24, 25].

Based on these findings, we tested the hypothesis that *Ca.* E. dacicola horizontal transfer can occur from a wild *B. oleae* population to adults of an artificially reared non-symbiotic colony. A second aim of this work was to find the easiest, fastest and most reliable method to detect *Ca.* E. dacicola DNA in *B. oleae* oesophageal bulb samples.

## Methods

**Insects** - Wild flies were obtained from infested olives harvested in several Tuscan olive orchards, during October - December 2015. Olives were kept in open boxes to maintain their freshness and to avoid fungi or mildew growth. A few days after harvesting, pupae were collected and transferred into plastic cages (BugDorm^®^, MegaView Science, Taiwan). Adults were supplied with sugar and water and kept at room temperature (18-20 °C).

Artificially reared *B. oleae* adults were obtained from a laboratory-adapted colony (Israel hybrid, IAEA, Seibersdorf, Vienna, Austria). Larvae were reared on a cellulose-based diet [26], while adults were reared in plastic cages (BugDorm^®^) and kept in a conditioned rearing room at 25°2 °C, RH 60°10%, and a 16:8 L:D photoperiod. Flies were supplied with water in a 30 mL plastic container with a sterile sponge strip acting as a wick and with a standard diet consisting of sugar, hydrolysed enzymatic yeast (ICN Biomedicals) and egg yolk (40:10:3).

**Experimental design** - Trials were started in February 2016. Since the goal was to transfer *Ca*. E. dacicola from a wild *B. oleae* population to a lab-reared one, the experiment was divided in two phases: a “contamination phase,” during which wild flies had time to contaminate different substrates, and an “acquisition phase,” in which lab flies were allowed to contact the substrates that had putatively been contaminated by *Ca.* E. dacicola. Before starting the experiment, the presence of *Ca.* E. dacicola in wild flies was confirmed by sequencing, as described below.

*Contamination phase* - Six treatments were tested as contamination sources: olives, gelled water, wax domes, wild faeces and cohabitation (lab females and wild males; lab males and wild females). The contamination sources are described below:

i. Olives - Freshly harvested ripe olives were given to 2-month-old wild adult flies to allow contamination with *Ca.* E. dacicola. Three Petri dishes with 30 olives each were put into a cage with more than 500 wild adults one week before the acquisition phase.
ii. Gelled water - Gelled water was given to 2-month-old wild adult flies to be contaminated by *Ca.* E. dacicola. Three Petri dishes with gelled water (8.35 g/L Gelcarin^®^, Duchefa Biochemie, The Netherlands) were put into a cage with more than 500 wild adults three days before the acquisition phase.
iii. Wax domes - Wax domes were used to collect eggs laid by wild flies; the domes were washed with a 2% sodium hypochlorite solution, rinsed twice in distilled sterile water and offered to 2-month-old wild adult flies to allow the females to oviposit. The resulting eggs were expected to be contaminated by *Ca.* E. dacicola based on previous research [27], and this was confirmed by sequencing. Three oviposition wax domes were placed into a cage with more than 500 adults two days before the acquisition phase.
iv. Wild faeces - Wild faeces were the fourth substrate used as a possible *Ca.* E. dacicola contamination source. One month before starting the acquisition phase, 100 wild flies ca. were put inside the cages assigned for the next phase (as described below) in order to contaminate the cage with their faeces.
v. Cohabitation between lab females × wild males - Cohabitation was used as a horizontal transfer method for *Ca.* E. dacicola, as described by Estes et al. [23]. The setup is described below.
vi. Cohabitation between lab males × wild females - The setup for this cohabitation method is described below.

*Acquisition phase* - Except for the faeces treatment, the next phase was started up in different cages (plastic boxes 2 L volume with a side closed by a nylon fine net, supplied with water and sugar) and set up as described below.

i. Olives - Three Petri dishes with olives putatively contaminated by *Ca.* E. dacicola were inserted into the plastic boxes (3 boxes, one dish each box) containing 25 male and 25 female newly emerged lab flies (younger than 24 h).
ii. Gelled water - Three Petri dishes with gelled water putatively contaminated by *Ca.* E. dacicola were inserted into plastic boxes (3 boxes, one dish each box) containing 25 male and 25 female newly emerged lab flies (younger than 24 h).
iii. Wax domes - Wax domes were opened and inserted on the bottom of the box (one each box) to let the lab flies get directly in contact with the eggs laid by the wild flies. The plastic boxes contained 25 male and 25 female newly emerged lab flies (younger than 24 h).
iv. Faeces - The 100 wild adults were removed from the dirty plastic boxes and 25 male and 25 female newly emerged flies (younger than 24 h) were transferred to each.
v. Cohabitation between lab females × wild males (labF × wildM) - Twenty-five newly emerged female flies (younger than 24 h) and 25 wild male flies of the same age were transferred into the plastic boxes.
vi. Cohabitation between lab males × wild females (labM × wildF) - Twenty-five newly emerged male lab flies (younger than 24 h) + 25 wild female flies of the same age were transferred into plastic boxes.

For each treatment, the acquisition phase lasted 15 days. Each treatment was replicated 3 times (6 trials with olives, gelled water, wax domes, faeces, labF × wildM, labM × wildF = 18 boxes, with a total of 900 tested flies). Boxes were arranged randomly on 4 shelves and moved daily to avoid any lighting bias. The setup of the overall experiment is summarized in Table 1.

**Table 1.**
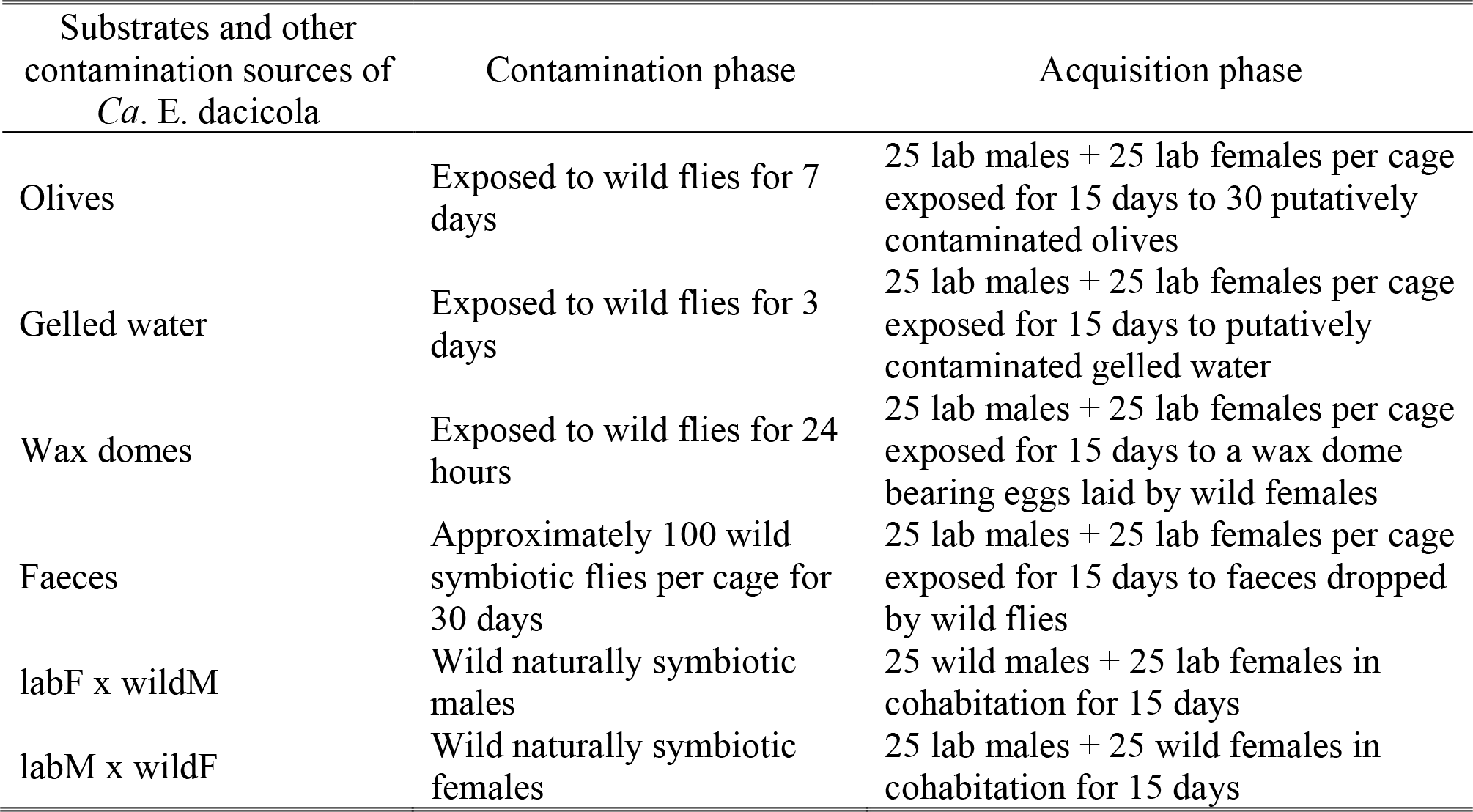
Setup of the horizontal transfer experiment

**Insect dissections** - After the acquisition phase, 30 flies were taken from each treatment (5 males and 5 females per cage for all three replicates), killed by freezing at −20 °C for 15 min and dissected. The dissection procedure was performed entirely under a laminar flow hood. Flies were first washed with a 2% sodium hypochlorite solution and then rinsed twice in distilled sterile water Second, each adult’s head was cut and opened under a stereoscopic microscope with sterile tools, and each oesophageal bulb was extracted. Sex, sample number and bulb aspect (transparent or milky) were noted. Finally, each bulb was put inside a 1.5 mL tube for DNA extraction.

**Culture-independent microbiological analyses** - Bacterial DNA from the oesophageal bulbs, faeces or sponge samples was extracted using 50 μL of InstaGene Matrix (BioRad Laboratories, Hertfordshire, UK) according to the manufacturer’s instructions. Bacterial DNA extracted from flies was obtained only from the oesophageal bulb and not from any other parts of the fly. Faeces were collected from the inner side of the cage top by rubbing sterile cotton on approximately 30 cm length. For the bacterial DNA extraction, the sterile cotton was treated as the oesophageal bulbs. Sponges were removed from the cages and transferred under laminar flow hood. Then, a small piece was removed with a scalpel and treated like the bulbs and faeces for the bacterial DNA extraction.

The extracted DNA was stored at −20 °C until PCR amplification. A preliminary PCR analysis was completed with EdF1 [15] and EdEnRev [10] primers designed to selectively amplify the 16S rRNA gene of *Ca.* E. dacicola. PCR-reactions were carried out using a T100 Thermal Cycler (BioRad Laboratories, Hertfordshire, UK) in 25 μl volumes containing 1X Flexi PCR buffer (Promega, Madison, WI), 1.5 mM MgCl_2_, 250 μM deoxynucleotide triphosphates (dNTPs), 400 nM of each primer, and 1U GoTaq^®^Flexi DNA polymerase (Promega). Amplifications were performed under the following conditions: an initial denaturation of 94 °C for 5 min followed by 35 cycles of 94 °C for 30 s, annealing at 55 °C for 30 s, extension at 72 °C for 45 s, and a final extension of 72 °C for 10 min. After PCR, the amplified products were verified by agarose gel electrophoresis (1.5% w/v), and the presumed presence/absence of *Ca.* E. dacicola in the specimens was scored based on the presence/absence of the targeted amplicon.

Additional primer sets were used in order to clarify the obtained results. For each primer set, the PCR reaction was carried out as described above. Ed1F was also paired with 1507R [28] to generate a nearly complete (1,300 bp) 16S rDNA fragment used for the subsequent screening of flies by ribosomal DNA restriction analysis (ARDRA). The 16S rDNA PCR products were digested separately with the restriction enzymes *Pst*I and *Cfo*I (Roche Diagnostics Ltd., Basel, Switzerland) as recommended by the manufacturer. The restriction fragments were separated by agarose gel electrophoresis (2% w/v), creating a specific restriction pattern for *Ca*. E. dacicola that distinguishes it from the other Enterobacteriaceae. The primer sets 986F-GC and 1401R [29] and 63F-GC and 518R [30] were used for the denaturing gradient gel electrophoresis (DGGE) analysis. PCR products were first verified by agarose gel electrophoresis (1.2% w/v) and successively loaded onto a polyacrylamide gel (40% acrylamide/bis 37.5:1; Serva Electrophoresis GmbH, Germany) containing a linear chemical denaturant gradient obtained with a 100% denaturant solution consisting of 40% v/v deionized formamide and 7 M urea. DGGE gels were run for 17 h at 60 °C and a constant voltage (75 V), using the Dcode DGGE System (Bio-Rad). After the electrophoresis gels were stained with SYBR^®^Gold (Molecular Probes, Eugene, OR) diluted 1:1,000 in 1X TAE buffer, the images were digitally captured under UV light (λ = 302 nm) using the ChemiDoc XRS apparatus (Bio-Rad). DGGE rDNA fragments from *Ca.* E. dacicola showed a distinct migration behaviour and could be easily distinguished from fragments derived from other oesophageal bulb-associated bacteria. PCR amplification and DGGE were also performed on DNA extracted from wild fly faeces and from sponges used as water wicks in each cage.

**Sequence analysis** - The middle portion of several DGGE-bands was aseptically excised and placed in 30 μL of distilled water. The PCR products were eluted from the gel through freezing and thawing and were subsequently re-amplified as described above and subjected to direct sequencing by Genechron (Ylichron, ENEA, Italy; http://www.genechron.it). Another subset of PCR products, obtained with the Ed1F and 1507R primers, was sequenced in both directions to verify the identity of *Ca.* E. dacicola in the oesophageal bulb specimens. The 16S rDNA sequence chromatograms were edited using Chromas Lite software (v2.1.1; Technelysium Pty, Ltd.http://www.technelysium.com.au/chromas-lite.htm) to verify the absence of ambiguous peaks and convert them to a FASTA format. The DECIPHER’s Find Chimera web tool (http://decipher.cec.wisc.edu) was used to uncover chimaeras hidden in the 16S rDNA sequences. The web-based BLAST tool available at the NCBI website (http://www.ncbi.nlm.nih.gov) was used to find taxonomically closely related nucleotide sequences. The nucleotide sequences identified in this study were deposited in the GenBank database under accession numbers MF095700 to MF095734.

## Results

**Screening** - As a result, PCR amplifications carried out with the primers EdF1 and EdEnRev highlighted a product with an expected size. A total of 17 of the 30 samples of wax domes, 26 of the 30 olive samples, 0 of the 30 gelled water samples, 16 of the 30 faeces treatment conditions, 16 of the 30 samples of labF × wildM, and 13 of the 30 samples of labM × wildF were found to be positive through PCR. As a double check, samples that were positive for the EdF1/EdEnRev amplification were screened by ARDRA. PCR products from both the wild flies and the cohabitation flies showed no recognition for the restriction enzyme *Pst*I; nevertheless, samples from lab reared flies and from those of other horizontal transfer crosses revealed the presence of one site for this enzyme (Figure 1), as previously described by Estes et al. [15]. ARDRA carried out with restriction enzyme *Cfo*I (Figure 2) revealed two unique patterns. One pattern corresponded to the wild fly samples and to those from the cohabitations, while the other pattern corresponded to the lab reared fly samples and those from the horizontal transfer cross. Bacterial DNA samples from oesophageal bulbs showing these two different patterns were re-amplified with EdF1/1507R primers and sequenced in both directions to obtain a nearly complete 16S rRNA gene sequence. Then, samples from wild flies, lab flies and from the horizontal transfer experiment crosses were tested by DGGE analysis, performed with the 986F-GC and 1401R primers. Visual inspection of DGGE revealed the presence of a single dominant band in all samples; in addition, some samples 11 also showed other less prominent bands (data not shown). Meanwhile, samples from wild flies and from most of the flies from cohabitations (n = 30) showed a similar migration pattern (data not shown). Despite this, the rest of the samples were found to have different fragment motilities. Successively, DGGE carried out with the 63F-GC and 518R primers was used to characterize the wild fly samples and compare them to those of the cohabitation fly samples. The DGGE profiles were comprised of a single dominant reoccurring band, as well as other less noticeable bands. All the profiles obtained from wild flies and most obtained from the cohabitation flies corresponded to one of the two main migration behaviours (Figure 3). A total of 6 unique bands separated by DGGE were selected according to their relative mobility, excised from the gel, and sequenced.

**Fig. 1.**
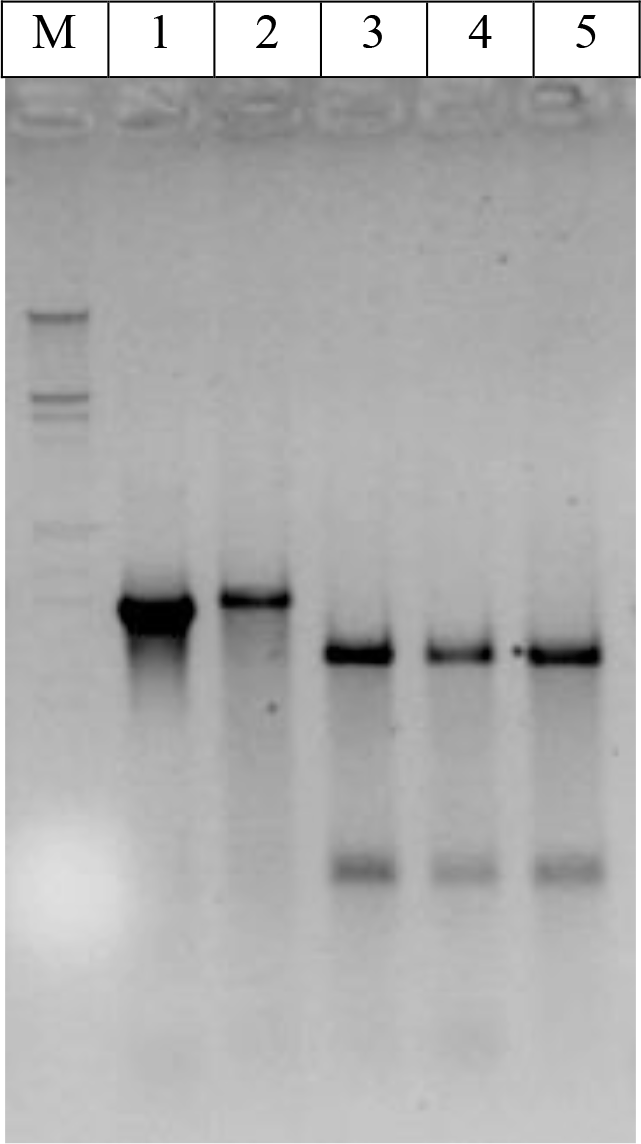
ARDRA patterns generated after the digestion of the amplified 16S rRNA gene with *Pst*I. Lane M corresponds to DNA Molecular Weight Marker III (Roche Diagnostics Ltd.), lane 1 corresponds to a non-digested 16S rDNA amplicon from a wild fly oesophageal bulb, lane 2 corresponds to the ARDRA pattern from a lab fly oesophageal bulb bacterial content, lane 3 corresponds to the ARDRA pattern from a wild fly oesophageal bulb bacterial content, and lanes 4 and 5 correspond to the ARDRA patterns from two lab fly oesophageal bulbs of the cohabitation treatment.

**Fig. 2.**
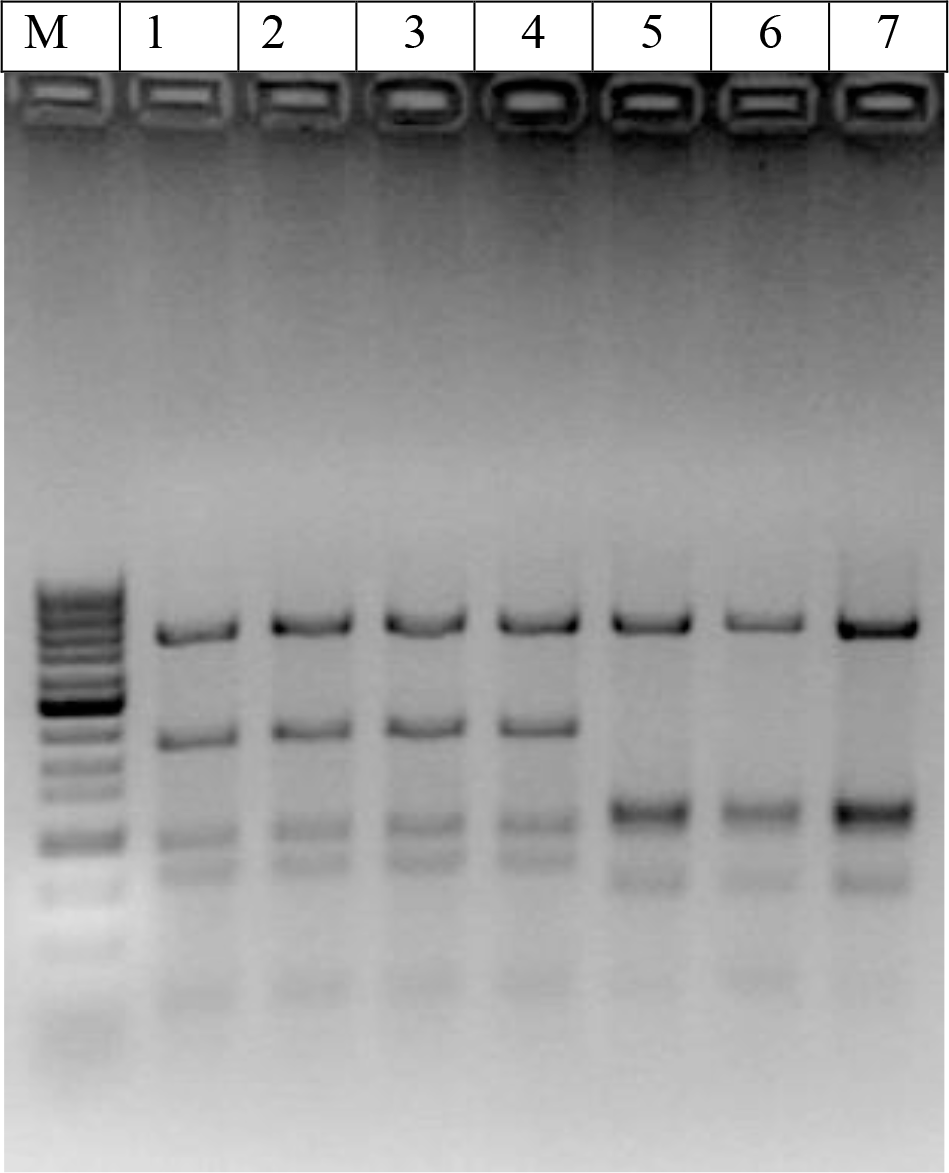
ARDRA patterns generated after digestion of the amplified 16S rRNA gene with *Cfo*I. Lane M corresponds to a 100 Base-Pair Ladder (GE Healthcare), lane 1 corresponds to the ARDRA pattern from a lab fly oesophageal bulb, lanes 2, 3 and 4 correspond to the ARDRA patterns of three lab fly oesophageal bulbs, lane 5 corresponds to the ARDRA pattern from a wild fly oesophageal bulb, and lanes 6 and 7 correspond to the ARDRA pattern from two lab fly oesophageal bulbs from the cohabitation treatment.

**Fig. 3.**
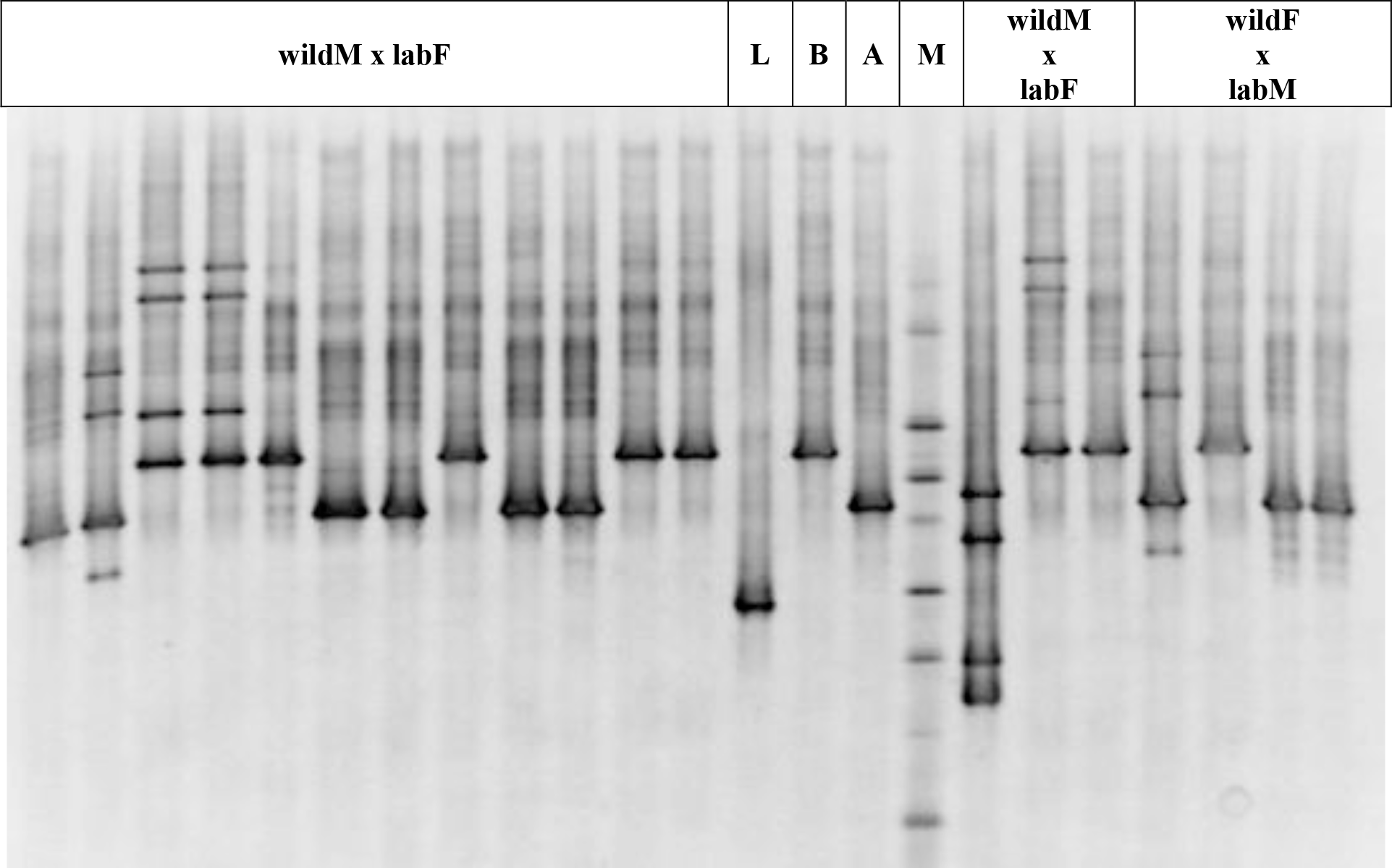
Analysis of the bacterial communities within the oesophageal bulbs of *B. oleae* after the cohabitation experiments: The DGGE profiles of 16S rDNA fragments obtained by amplification with the 63FGC/518R primer set. The letter M on the gel image indicates the marker used for the normalization of the bands in the profiles. L refers to a lab sample, while B and A correspond to the two different *Ca*. E. dacicola lineages from wild flies (htB and htA, respectively). The other headings refer to the two different cohabitation treatments.

**Sequencing** - The presence of *Ca. E*. dacicola in the oesophageal bulb samples of wild flies was confirmed before starting the horizontal transfer experiment by sequencing the PCR products (n = 6) obtained using EdF1 and 1507R primers. In all cases, we obtained species-level identity ascribed to the sequence of *Ca.* E. dacicola (100% similarity to GenBank accession number HQ667589 or HQ667588). PCR products (n = 3) amplified from the oesophageal bulbs of lab-reared flies were also sequenced to obtain species-level identity with the sequence of *Morganella morganii* (99% similarity to GenBank accession number NR_113580). By sequencing, the DGGE isolate (n = 2) bands of the wild fly specimens were confirmed to correspond to the sequence of *Ca.* E. dacicola (>99% similarity). In particular, the lower band (Figure 3) was assigned to *Ca.* E. dacicola haplotype A (GenBank accession number HQ667588) and the upper band (Figure 3) to *Ca.* E. dacicola haplotype B (GenBank accession number HQ667589), as already distinguished by Savio et al. [24]. The exclusive incidence of *Ca.* E. dacicola was additionally confirmed in 4 isolated DGGE bands of fly specimens from the cohabitation experiments, which demonstrated similar migration behaviours to the wild fly samples. On the other hand, the DGGE isolate (Figure 3) bands of the lab-reared flies were found to share sequence identity with *M. morganii* (99% similarity to GenBank accession number NR_043751). Other bands showing different migration behaviours from those of the wild or lab flies were not sequenced.

**Faeces and Sponges** - PCR-DGGE analyses of wild flies’ faeces (Figure 4) and the subsequent sequencing of the excised DGGE bands provided evidence of the presence of taxa mainly related to the γ-Proteobacteria phylum and, in particular, to the Enterobacteriales order (Table 2). The nucleotide-sequence identities ranged from 91% to 100%, and most matches showed identities greater than 99%. *Ca.* E. dacicola was also found (with 100% similarity to GenBank accession number HQ667589), although it was detected as a less pronounced band and a narrow denaturing gradient needed to be applied to highlight its presence in the faeces samples (Figure 4B). Furthermore, PCR-DGGE analyses performed on the sponges highlighted the presence of *Ca.* E. dacicola on those taken from the replicates of the faeces treatment (data not shown). Analyses on the sponges from different treatments (olives, wax domes, cohabitation and gelled water cages) did not show any match with the *B. oleae* endosymbiont.

**Fig. 4.**
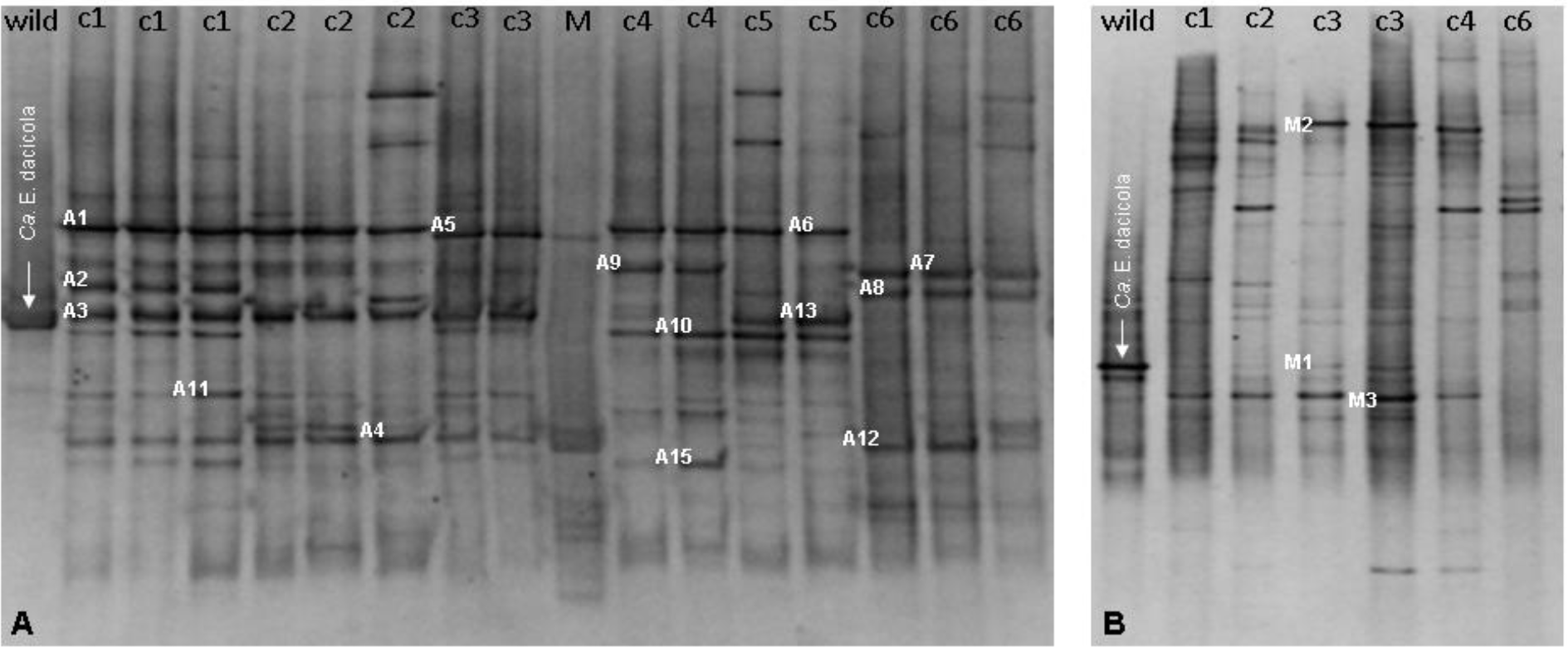
Analysis of the bacterial communities within the faeces of *B. oleae:* DGGE profiles of the 16S rDNA fragments obtained by amplification with the 986FGC/1401R primer set. DGGE denaturing gradients of 45-68% (A) and 50-65% (B). The arrowed bands indicate the PCR products obtained by the amplification of DNA extracted from the wild fly oesophageal bulbs used as species markers of *Ca.* E. dacicola. Numbered bands (A1-A15; M1-M3) were selected for sequencing. The faeces were deposited by wild fly samples in cages 1-5 (c1-c5) and by lab flies in cage 6 (c6), with 2 or 3 replicates for each cage. M, marker.

**Table 2.**
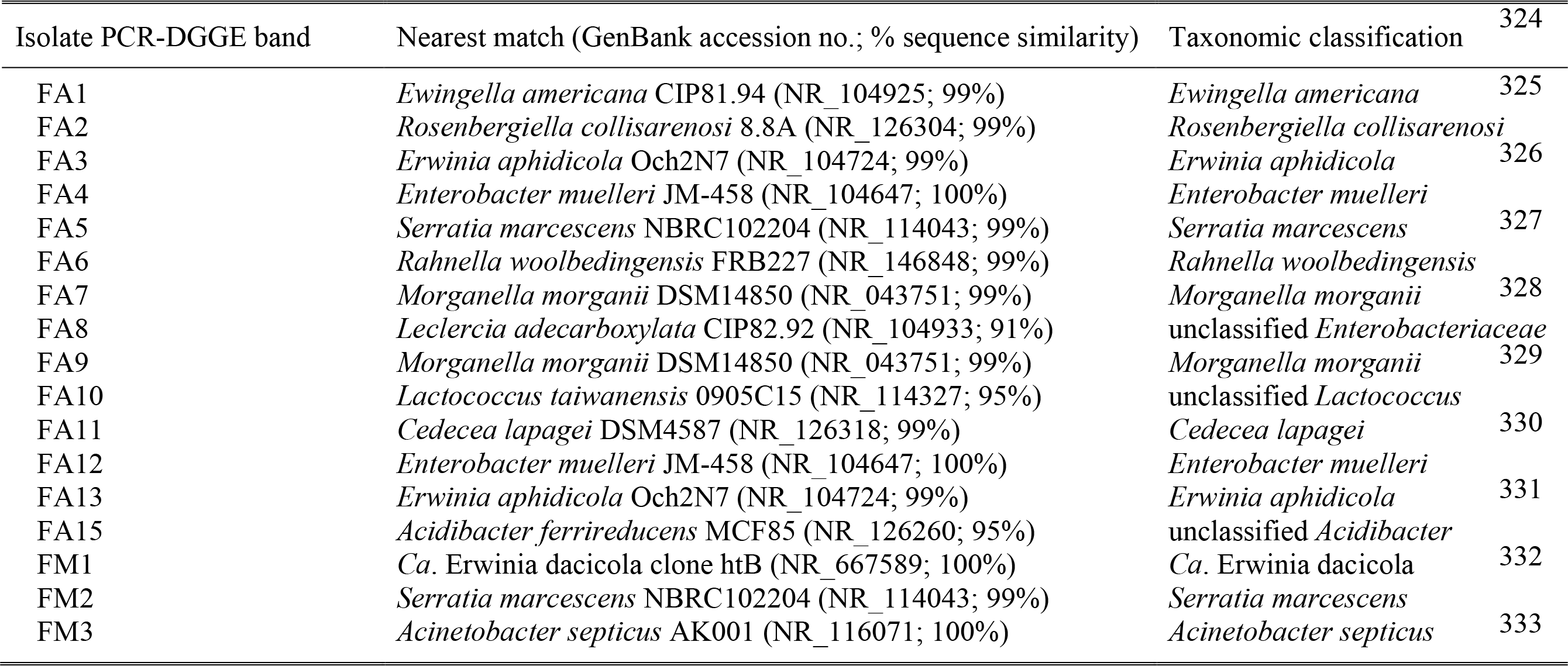
Identification of 16S rDNA fragments selected from PCR-DGGE of the *B. oleae* faeces. Taxonomic identification was achieved using different sequence similarity thresholds: a similarity ≥97% was used for species level identification, while similarities of 95%, 90%, 85%, 80% and 75% were used for assignment at the genus, family, order, class and phylum levels, respectively [31].

## Discussion

The goal of these investigations was to attempt to observe the horizontal transfer of the endosymbiont *Ca.* E. dacicola from a wild *B. oleae* population to a laboratory colony. A secondary goal was to determine the best and most efficient method to reliably screen for this endosymbiont in *B. oleae* samples. It was predicted that horizontal transfer could occur via both oral contamination (wild flies’ regurgitation on gelled water and olives) and via anus-genital contamination (eggs laid inside oviposition domes by wild flies, wild faeces, and cohabitation with wild flies).

Concerning the oral contamination transmission route and more specifically regurgitation, we tested the hypothesis that flies could regurgitate saliva with bacteria on two different substrates, olives and gelled water. Petri first described this behaviour in 1907 [32], and he reported a peculiar behaviour of *B. oleae* in which the fly sucked and regurgitated olive juice during the oviposition process, commonly known as “the kiss” [33]. Tzanakakis [34] also described this action in *B. oleae*, assuming that, at the end of the oviposition process, the female retracts the ovipositor and regurgitates the juice sucked from the hole to deter subsequent oviposition. Drew and Lloyd [35] also described strict relationships between tropical Dacinae and the bacteria of host plants. They showed that the bacteria present in the alimentary tract of flies were also found on the surface of host fruit from plants in which flies had been collected, suggesting that regurgitation was involved in this bacterial presence. However, in our experiment, even if the substrates had been contaminated through bacterial regurgitation by the wild olive fruit fly, the transfer of *Ca.* E. dacicola to lab flies did not occur, either through the olives or the gelled water. However, no attempts to detect *Ca.* E. dacicola on these two substrates were carried out, since the transfer did not occur we presume that the symbiont was not present on them or, if present, it was probably not available for the horizontal transfer.

Regarding the possible anus-genital transfer, wax domes containing eggs laid by wild females were tested as a contamination source. The presence of *Ca.* E. dacicola was found on the eggs, not only by biological molecular techniques [10] but also by morphological observations dealing with the presence of bacterial colonies around the ano-genital opening and in the micropylar area [6]. Furthermore, previous observations had highlighted the presence of bacterial masses on *B. oleae* eggs [36]. Since several previous studies demonstrated that *Ca.* E. dacicola is vertically transmitted from the female to the egg [9, 10, 15, 17, 25]; we predicted that a horizontal transfer mechanism could occur after the lab flies have direct contact with the eggs laid by wild females. However, our attempt was not successful. In terms of vertical transmission, there are many ways to “pass” symbiotically useful bacteria via the egg, from the mother to the progeny. For instance, symbiotic bacteria can be maternally transmitted by “capsule transmission” or by “egg smearing,” as observed in stinkbugs [37]. It could also be transferred to the egg as it passes through the micropyles, as is believed to occur in fruit flies [38]. For the vertical transfer of *Ca.* E. dacicola in *B. oleae*, the bacterium seems to be maternally transmitted by “egg smearing” [6]. Thus, even if the endosymbiont is smeared on the egg’s surface, its passage to the young larva is probably strictly related to the micro environment inside the olive. Given these assumptions, we predict that in the present work, this horizontal transfer via egg using wax domes did not occur, perhaps because *Ca.* E. dacicola on the egg surfaces was exposed to air for too long, instead of remaining in the “small oblong chamber” inside the olive [34] with low oxygen levels, thus limiting the possibility of horizontal transfer. Another hypothesis could be that after oviposition inside the fruit, the endosymbiont needs some olive compounds that enable it to stay viable until larval assumption. Because the symbiont passes through and colonizes the digestive tract during the entire adult lifespan [15], and especially given its role in nitrogen metabolism [25], we tested the hypothesis that it could be partially released in the faeces after digestion. The endosymbiont was indeed detected on faeces and on sponges taken from the replicates of the faeces treatment. These sponges stayed in contact with the wild flies for a long time (they were inserted during the contamination phase along with wild adults, and they were not exchanged with new sterile sponges for the acquisition phase, as in other theses). We therefore believe that they were contaminated by faeces.

However, no horizontal transfer was observed after using this substrate as a contamination source. Based on this, we presume that even if *Ca.* E. dacicola DNA was detected both on the faeces and sponges, the bacterium may not be viable or may on these substrates and may not be horizontally transferred in this way. These findings further suggest that *Ca.* E. dacicola may be a bacterium that needs low levels of oxygen to maintain its vitality and grow.

Consistent with our hypotheses and the results of Estes et al. [23], horizontal transfer via cohabitation with wild flies was the only treatment in which transfer occurred. To our knowledge, the transmission of *Ca.* E. dacicola could have occurred through different methods, including mating, coprophagy or trophallaxis. Copulation between males and females was not directly verified; there is a high probability that the flies did mate, but we cannot be sure that this was the way through which the transfer occurred. Further trials assessing cohabitation between wildM × labM or wildF × labF could be set out in order to better clarify this finding. The flies in the cohabitation scenario also had ample opportunities to regurgitate and defecate in the same cage. This observation allowed us to make a second hypothesis: perhaps not only the mating, but also the coprophagy and/or the trophallaxis behaviour between wild and lab flies during their cohabitation accounted for the horizontal transfer. The only thing we know is that the wild and lab flies stayed together for 15 days and they had time to perform other behaviours and to be in contact frequently in different ways. Trophallaxis represents an “exchange of alimentary liquid among colony members and guest organisms,” and it can occur before, during, or after mating. It can also be direct or indirect, stomodeal or proctodaeal, and it has been described in approximately 20 species of Tephritidae, representing a behaviour that involves the transfer of substances [39]. Several studies described the mating trophallaxis in Tephritidae [40, 41, 42] but did not demonstrate the transfer of any substance during the contact between the mouthparts of the mates. Our results lead us to suppose that this behaviour could be involved in endosymbiont transfer, as predicted by Estes et al. [23]. They hypothesized that bacterial transfer occurs through coprophagy, presumably thanks to pre/in direct proctodaeal trophallaxis. Moreover, it must be noted that we found *Ca.* E. dacicola

DNA inside the oesophageal bulb of lab flies that cohabited with wild flies; as a consequence, trophallaxis appears to be more likely to be responsible for transfer than *Ca.* E. dacicola matings. Further research, such as the analysis of the proctodaeal diverticula and/or the crop system of lab flies after cohabitation with wild adults, together with behavioural studies, would better clarify this aspect. Moreover, cohabitation was the only treatment in which the endosymbiont was not as exposed to oxygen. In contrast, the other treatment conditions, such as the olives, gelled water, eggs laid by wild females and faeces likely exposed to *Ca.* E. dacicola, were all exposed to oxygen for a longer period. We can therefore presume that *Ca.* E. dacicola prefers microaerophilic conditions for its vitality and transfer. In addition, we can affirm that transfer via cohabitation is not related to the sex of the wild symbiotic fly, since it occurred both when the *Ca.* E. dacicola contamination sources were wild females or wild males.

Hence, a symbiotic wild fly (male or female) in cohabitation with a non-symbiotic lab fly (male or female) is all that is required for the successful horizontal transfer of *Ca.* E. dacicola. Thus, this could be the first step in obtaining a permanently symbiotic laboratory olive fruit fly colony, likely reared on different substrates than the cellulose-based one, which allow for the avoidance of genetic modifications possibly caused by symbiont absence [19, 20].

The aim of the present study was to provide a reliable and consistent tool for implementing the detection of the endosymbiont in a large number of *B. oleae* specimens and/or environmental samples. According to the obtained results, it seems that the primers EdF1 and EdEnRev are not sufficiently specific for *Ca.* E. dacicola, as previously described by Estes *et al.* [15]. Indeed, samples that were positive to *Ca.* E. dacicola with these primers did not show the same results after DGGE analysis. Moreover, an *in silico* analysis conducted using the Probe Match function within the RDP-II database (http://rdp.cme.msu.edu) showed a higher number of exact matches to the 16S rRNA gene sequences from members of Enterobacteriaceae family (3% respect to the total Enterobacteriaceae sequences in RDP database) belonging to *Erwinia, Serratia, Proteus,Buttiauxella, Enterobacter* and other genera. Thus, we suggest that to confirm the presence of *Ca.* E. dacicola, the screening of oesophageal bulbs or other specimens by PCR with EdF1/EdEnRev primer has to be combined with subsequent analyses [27]. Sequencing is a time consuming and expensive method, and this does not seem to be the most convenient system, especially when a large number of samples must be analysed. ARDRA has been previously and successfully performed to compare profiles from uncultivable bulk bacteria residing in the oesophageal bulb with those from cultivable bacteria occasionally arising on plates in an attempt at endosymbiont isolation (Capuzzo et al., 2005) and, more recently, to distinguish the two different bacterial haplotypes (htA and htB) [24]. Furthermore, Ben-Yosef *et al.* [25] used DGGE performed with 986F-1401R primers and succeeded in detecting *Ca.* E. dacicola in *B. oleae* adult oesophageal bulbs and larvae. In this study, both ARDRA and DGGE techniques were applied. ARDRA demonstrated that it was possible to identify a specific profile corresponding to *Ca.* E. dacicola that was clearly distinguishable from that of other Enterobacteriaceae, such as *M. morganii.* Moreover, DGGE appears to be the best molecular fingerprinting method, since different bacterial taxa may be associated with oesophageal bulbs, both as individual dominant bacterium and in the bacterial consortium. The PCR-DGGE fingerprint was widely used to compare the microbial community structure in a variety of environments [43, 44, 45, 46]. Furthermore, it supports the identification of bands, because PCR products can be recovered and sequenced [47]. As an alternative to sequencing, the identification of bacteria may be achieved by the comparison of the PCR amplicon DGGE migration behaviour with that of a reference strain, used as species marker [48]. Thus, the choice of which target hypervariable regions of the 16S rRNA gene are to be amplified may strongly affect the quality of information obtained by DGGE [47]. This study demonstrated that PCR-DGGE performed with the primer set 63F-GC/518R and targeting the V1-V3 hypervariable regions, provides the best procedure for the rapid and straightforward screening of the presence of *Ca.* E. dacicola in a high number of fly specimens. This also reflects the two different *Ca.* E. dacicola haplotypes (htA and htB).

Considering the ARDRA profiles and the migration behaviour of PCR products on DGGE and nucleotide-sequence identity by BLAST, approximately 50% of the oesophageal bulbs of lab flies after cohabitation highlighted the presence of *Ca.* E. dacicola as a prominent associated species, and in particular, 13 corresponded to *Ca.* E. dacicola haplotype A and 13 to *Ca.* E. dacicola haplotype B, confirming previous findings from fly samples collected in Tuscany [24]. Conversely, all the oesophageal bulbs of the lab-reared flies of the other crosses in the horizontal transfer experiment did not demonstrate the acquisition of *Ca.* E. dacicola. Furthermore, the other associated bacteria were supposed to be related to different taxa within the Enterobacteriaceae family.

The fact that *M. morganii* was detected in lab flies shows that the lab strain has been exposed to many bacteria and that *M. morganii* could have competed with *Ca*. E. dacicola, thus preventing horizontal transfer. This does not mean that *M. morganii* could represent a pathogen for *B. oleae*, as shown in recent studies on *Anastrepha* spp. [49, 50]. Furthermore, this bacterium has already been found in the oesophageal bulb of lab-reared *B. oleae*’s flies [13] and does not seem to represent a threat for the olive fruit fly. Along with this, supplementary observations would be appropriate to better evaluate the effects of this bacterium on *B. oleae* fitness and other parameters such as adult mortality or egg production.

## Conclusions

This research demonstrates that the cohabitation of wild and lab reared flies is the only way through which the horizontal transfer can occur. Thanks to these investigations, it has been possible to find a viable way to transfer the endosymbiont *Ca.* E. dacicola from an adult wild *B. oleae* population to a laboratory colony. As a result, this study represents the first step in better understanding *Ca.* E. dacicola behaviour, physiology and culturing requirements.

DGGE was the most reliable detection method, although it has some inherent associated limitations; DGGE proved to be a consistent method for screening the endosymbiont *Ca.* E. dacicola in *B. oleae*, further distinguishing between the two *Ca.* E. dacicola haplotypes.

Further investigations should be completed in order to improve these findings, and other horizontal transfer experiments should be completed during different periods of the year and/or in different conditions. Moreover, the resulting endosymbiotic laboratory-reared flies should be evaluated in terms of different parameters, such as egg production, egg hatching, larval development and pupal recovery for the pre-imaginal stages and mortality, lek behaviour and mating success for the adult stages. Nevertheless, the trials in which the transfer did not occur (olives, gelled water, wax domes, faeces) may be tested again using a different approach to better understand how to solve the problems that hindered the transfer. In this way, different strategies could be identified in order to improve the success of the horizontal transfer. Thus, laboratory-reared flies could compete with the wild ones, improving the Sterile Insect Technique as a possible tool for the sustainable control strategies within the olive system.

## Author contributions

All authors conceived of and designed the experiments. GB and RG reared the insects and performed the experiments. GB and PS performed insect dissection and DNA extraction. RP designed the molecular biology procedures. GB and RP performed the PCR analyses, DGGE, ARDRA and other molecular techniques applied to the samples. GB and RP drafted the manuscript; AB and PS discussed and revised the initial draft of the manuscript.

All authors read and approved the final version of the manuscript.

## Acknowledgements

The authors are grateful to Carlos Càceres (Insect Pest Control, IAEA, Seibersdorf, Austria) for supplying the starting colony of Israel hybrid *B. oleae.* Antonio Belcari and Patrizia Sacchetti also acknowledge the International Atomic Energy Agency of Vienna, Austria, for supporting their participation in the CRP “Use of Symbiotic Bacteria to Reduce Mass-Rearing Costs and Increase Mating Success in Selected Fruit Pests in Support of SIT Application” (D41024). This research was partially funded by the University of Florence (Fondi d’Ateneo).

